# Pathogenic soluble tau peptide disrupts endothelial calcium signaling and vasodilation in the brain microvasculature

**DOI:** 10.1101/2023.08.08.552492

**Authors:** Kalev Freeman, Adrian M Sackheim, Amreen Mughal, Masayo Koide, Grace Bonson, Grace Ebner, Grant Hennig, Warren Lockette, Mark T Nelson

## Abstract

The accumulation of the microtubule-associated tau protein in and around blood vessels contributes to brain microvascular dysfunction through mechanisms that are incompletely understood. Delivery of nutrients to active neurons in the brain relies on capillary inositol 1,4,5-triphosphate receptor (IP_3_R)–mediated calcium (Ca^2+^) signals to direct blood flow. The initiation and amplification of endothelial cell IP_3_R-mediated Ca^2+^ signals requires an intact microtubule cytoskeleton. Since tau accumulation in endothelial cells disrupts native microtubule stability, we reasoned that tau-induced microtubule destabilization would impair endothelial IP_3_-evoked Ca^2+^ signaling. We tested the hypothesis that tau disrupts the regulation of local cerebral blood flow by reducing endothelial cell Ca^2+^ signals and endothelial-dependent vasodilation. We used a pathogenic soluble tau peptide (T-peptide) model of tau aggregation and mice with genetically encoded endothelial Ca^2+^ sensors to measure cerebrovascular endothelial responses to tau exposure. T-peptide significantly attenuated endothelial Ca^2+^ activity and cortical capillary blood flow *in vivo* within 120 seconds. Further, T-peptide application constricted pressurized cerebral arteries and inhibited endothelium-dependent vasodilation. This study demonstrates that pathogenic tau alters cerebrovascular function through direct attenuation of endothelial Ca^2+^ signaling and endothelium-dependent vasodilation.

## INTRODUCTION

Impaired cerebral blood flow plays a crucial role in the development of degenerative neurological diseases such as Alzheimer’s disease (AD) and Chronic Traumatic Encephalopathy (CTE). Both AD and CTE are characterized by accumulation of hyperphosphorylated forms of the microtubule-associated protein tau around blood vessels ^1–5^. Tau disrupts cerebral blood flow regulation, but mechanisms are incompletely understood ^6,7^. The precise and efficient delivery of nutrients to active neurons in the brain relies on the intricate network of capillaries in which local inositol 1,4,5-triphosphate IP_3_ receptor (IP_3_R)–mediated calcium (Ca^2+^) signals through nitric oxide production direct blood flow ^8^. These events range from small, brief proto-events, which represent the release of Ca^2+^ through a few IP_3_Rs channels, to larger, sustained compound events lasting up to approximately one minute, involving clusters of channels ^8^. As a result, intracellular Ca^2+^ activates nitric oxide synthesis, which enhances local blood flow to active neurons.

It was recently demonstrated that brain microvascular endothelial cells internalize soluble tau oligomers in pathological conditions through a process that depends on heparan sulfate proteoglycans ^7^. Endothelial cell internalization of soluble tau triggers phosphorylation of endogenous tau at threonine 231, which promotes destabilization of microtubules and reduces microtubule density ^7^. This has important implications, because microtubule function is required for IP_3_R clustering and Ca^2+^ signaling in endothelial cells. Prior work has shown that perturbation of the microtubule cytoskeleton by the drugs colchicine and colcemid prevents the initiation of IP_3_-evoked Ca^2+^ signals in endothelial cells ^9,10^. Furthermore, disruption of microtubules with nocodazole impairs the microtubule-facilitated motility of IP_3_Rs, preventing the localization of the channel which is needed for efficient and graded recruitment of channels with increasing stimulus strength ^11^. Microtubules also have direct interactions with the consensus motif for EB binding (TxIP) of IP_3_Rs through the microtubule end-binding protein 3 (EB3); depletion or deletion of EB3, or mutation of the TxIP motif on IP_3_Rs, prevents clustering and functional coupling between IP_3_Rs ^12^. Since tau accumulation in endothelial cells disrupts native microtubule stability, we reasoned that tau-induced endothelial microtubule destabilization would impair endothelial IP_3_-evoked Ca^2+^ signaling.

Here, we tested the hypothesis that tau aggregates impair the regulation of regional cerebral blood flow by acutely reducing endothelial cell Ca^2+^ signals and disrupting endothelium-dependent vasodilation. We used a soluble pathogenic tau peptide (T-peptide) model of tau aggregation. This cell-permeable hexameric peptide derived from the third microtubule-binding repeat of tau self-assembles *in vitro* into paired helical filament-like aggregates providing an established paradigm for studying mechanisms underlying tauopathy ^13,14^. To isolate and measure Ca^2+^ activity from endothelial cells, we studied mice with endothelial-specific expression of the Ca^2+^ biosensor GCamP8 driven by the cadherin 5 promotor (GCaMP8-*Cdh5* mice) ^8^. We utilized two-photon laser scanning microscopy via a cranial window to record microvascular endothelial Ca^2+^ signals *in vivo*; the addition of intravenous FITC-dextran allowed measurement of regional cerebral blood flow. We also isolated small cerebral arteries to study direct vascular effects of T-peptide with pressure myography. Together, these experiments demonstrate that T-peptide has rapid and direct effects on brain microvascular endothelium, reducing Ca^2+^ signaling and impairing the regional regulation of cerebral blood flow.

## METHODS

### Animals

Male C57BL/6j mice (Jackson Laboratories, Bar Harbor, ME), aged 8-12 weeks, were assigned at random to either experimental or control treatment groups. *Cdh5*-GCaMP8 mice (male, age 8-12 weeks) were used for *in vivo* Ca^2+^ imaging studies. Mice were kept on a 12 h light/dark cycle, with *ad libitum* access to food and water and were housed in groups of five. All studies were conducted in accordance with the Guidelines for the Care and Use of Laboratory Animals (National Institute of Health), Guidelines for Survival Rodent Surgery (National Institute of Health) and approved by the Institutional Animal Care and Use Committee of the University of Vermont. All procedures were conducted and reported in accordance with the ARRIVE guidelines (Animal Research: Reporting In Vivo Experiments) (https://www.nc3rs.org.uk/arrive-guidelines).

### In vivo imaging of cerebral hemodynamics

Mice were anesthetized with isoflurane (5% induction; 2% maintenance). The skull was exposed, a stainless-steel head plate was attached, and a small circular cranial window was created over the somatosensory cortex. Approximately 150 μL of a 3-mg/mL solution of FITC or TRITC-dextran (MW 150 kDa) in sterile saline was injected intravenously into the retro-orbital sinus to allow visualization of the cerebral vasculature and contrast imaging of red blood cells (RBCs). Upon conclusion of surgery, isoflurane anesthesia was replaced with α-chloralose (50 mg/kg) and urethane (750 mg/kg). Body temperature was maintained at 37°C throughout the experiment using an electric heating pad. The brain was continuously perfused with artificial cerebrospinal fluid (aCSF) containing (in mM): 124 NaCl, 3 KCl, 2 CaCl_2_, 2 MgCl_2_, 1.25 NaH_2_PO_4_, 26 NaHCO_3_, and 4 glucose (pH 7.4). A pipette was introduced into the cortex, maneuvered adjacent to the capillary under study and pressurized injections (200–300 ms, 7 ± 1 PSI) of 100 nM T-peptide (Tocris, USA) with tetramethylrhodamine isothiocyanate (TRITC, MW 150 kDa; 0.06 mg/mL)-labeled dextran were ejected directly onto the capillary. RBC flux data was collected by line scanning the capillary of interest at 15 kHz. Single-plane imaging experiments involved imaging a 425 µm x 425 µm field at ∼2.5 Hz for 360 s. Images were acquired through a Zeiss 20x Plan Apochromat 1.0 NA DIC VIS-IR water-immersion objective mounted on a Zeiss LSM-7 multiphoton microscope (Zeiss, USA) coupled to a Coherent Chameleon Vision II Titanium-Sapphire pulsed infrared laser (Coherent, USA). *Cdh5*-GCaMP8 and TRITC-dextran were excited at 920 nm, and emitted fluorescence was separated through 500- to 550- and 570- to 610-nm bandpass filters, respectively.

### Ex vivo Pressure Myography

Cerebral vessels were harvested and pressurized in a myograph system as previously described ^15^. Briefly, mice were euthanized and a full craniectomy was performed to expose the brain, which was then dissected and placed into cold (4°C) HEPES-buffered physiological saline solution (HEPES-PSS) with the following composition (in mM): 134 NaCl, 6 KCl, 2 mM CaCl_2_, 1 MgCl_2_, and 7 glucose (pH 7.4). The posterior cerebral artery (PCA) was then excised and cannulated in a pressure myograph chamber (Living Systems Instrumentation, USA) as previously described.^15^ Briefly, arteries were maintained in aCSF solution at 37° C and allowed to equilibrate for 10 minutes at 10 mm Hg before experimentation. Intraluminal pressure was controlled using a pressure servo system (Living Systems Instrumentation, USA) and blood vessel diameters were continuously recorded using edge-detection software (IonOptix, USA). Intraluminal pressure was increased to 80 mm Hg and vessels were allowed to develop spontaneous myogenic tone.

Endothelial viability was tested using 1 µM NS309 (Cayman Chemical, USA) and a dilatory response > 85 % was considered “intact.” Experimental interventions were then carried out using T-peptide (20 minutes; 100 nM), bradykinin (10 µM; Cayman Chemical, USA), NS309 (1 µM), or spermine NONOate (1 – 100 nM; Cayman Chemical, USA). The bath solution was then exchanged with 0-Ca^2+^ saline solution with 100 µM diltiazem (Sigma-Aldrich, USA) to elicit maximal passive diameter. The percentage change in diameter to pharmacological intervention was measured at the point of peak dilation or constriction and calculated as: change in diameter (%) = [(diameter after drug administration – baseline diameter)/(passive diameter – baseline diameter) x 100].

### Data Analysis & Statistics

GraphPad Prism software (ver. 9.4.1; GraphPad Software, USA) was used for X–Y graphing and analysis; values are presented as means ± standard deviation (SD). Data were tested for normality and appropriate parametric or non-parametric statistical tests were subsequently applied. Differences were considered significant if P <0.05. RBC flux was analyzed offline using custom software (SparkAn, Adrian Bonev, University of Vermont, USA). Flux data were binned at 1-second intervals. Mean baseline flux data for summary figures were obtained by averaging the baseline (6 second) for each measurement before pressure ejection of 100 nM T-peptide. The peak response was defined as the peak 1-s flux bin after delivery of T-peptide within the remaining scanning (240-300 seconds). The depth of capillaries below the surface was estimated from z-stack series acquired before pipette placement. To avoid limitations of standard F/F_0_ methods and accurately characterize and analyze Ca^2+^ events in brain capillaries, we used our custom autonomous image analysis program ‘VolumetryG9’ ^8^. This method uses signal-to-noise ratios and statistical z-scores (Zscrs) combined with spatiotemporal (ST) mapping to accurately quantify the full dimensional characteristics of Ca^2+^ events, combining amplitude (Zscr), size (µm) and duration (s) as the conglomerate variable ZUMS (**Z**scr·**μm**·**s**econd) (8). Each animal or vessel served as its own control and experiments were analyzed in a pairwise fashion; therefore anonymizing was not feasible. Sample sizes for all experimental preparations were estimated using our previous experience ^16^.

## RESULTS

### Cortical capillary blood flow and EC Ca^2+^ signaling are significantly reduced by T-peptide *in vivo*

Targeted delivery of nutrients to active neurons in the brain depends on capillary endothelial cell (IP_3_R)–mediated Ca^2+^ signals. Since tau accumulation in microvascular brain endothelial cells disrupts the native microtubule function ^7^, and these microtubules are needed for initiation and amplification of the IP_3_R-mediated Ca^2+^ signals, we postulated that tau exposure would lead to a rapid reduction in endothelial Ca^2+^ signaling. To test this hypothesis, we leveraged our recently developed *Cdh5*-GCaMP8 transgenic mice that express the Ca^2+^ indicator, GCaMP8, under the transcriptional control of the *Cdh5* (cadherin 5) promoter for targeted expression in vascular endothelial cells ^8^. For these experiments, we injected TRITC-dextran and used multi-photon microscopy to visualize brain capillaries through a cranial window over the somatosensory cortex. This provided a map of the cortical vasculature using 3-dimension z-stacks in a 425 x 425 µm field of view before and after low pressure ejection of T-peptide (100 nM; 300 msec; 6-8 PSI) on the cortical surface. We defined capillaries as vessels located between penetrating arterioles and ascending venule and assigned them branch orders as previously described ^8,17^. We then recorded the capillary endothelial cells (cECs) Ca^2+^ activity in layer II-III of the somatosensory cortex before and after cortical application of T-peptide (100 nM) (Fig1, A-D).

Under baseline conditions, both groups of mice demonstrated endothelial Ca^2+^ activity throughout the visual field. Within 2 minutes (120 seconds) of T-peptide exposure, we observed a significant reduction in Ca^2+^ activity from baseline (39% reduction; 8423 ± 2284 vs. 3257 ± 1130 ZUMS; n=5; P<0.05) (Figure 1 A-D). Of note, T-peptide did not reduce the number overall active sites, determined by the count of Ca^2+^ events throughout the visual field during the recording period (# events, 164 ± 103 vs. 104 ± 48; n=5; n.s.). This indicates that T-peptide decreases the Ca^2+^ output generated from individual active sites without impacting the number of sites themselves. This observation is consistent with a mechanism in which T-peptide disrupts microtubule function needed for IP_3_R clustering and efficient intracellular Ca^2+^ release. T-peptide exposure also resulted in a decrease in regional blood flow in 3^rd^-4^th^ order branches capillaries (Figure 1 E-H). We observed a rapid reduction red blood cell (RBC) flux within 2 minutes after T-peptide exposure compared to baseline, that persisted for the duration of the 5-minute recording period (34 ± 17 vs. 11 ± 8 RBCs/s; n=7; P<0.05).

**Figure 1.**
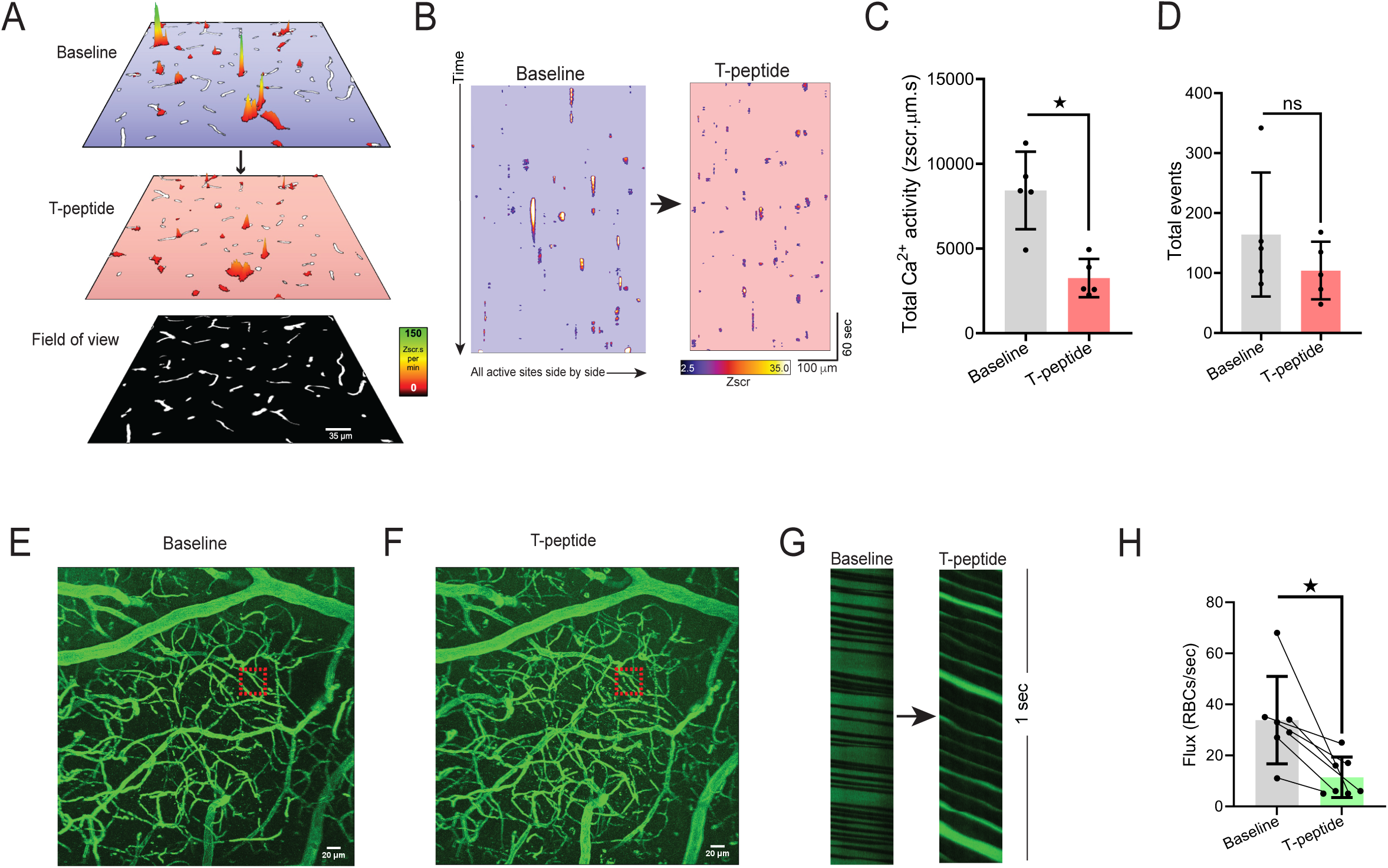
T-peptide significantly diminishes total Ca^2+^ activity in capillaries and regional RBC flux *in vivo.* (A) Prevalence maps from the same field of view showing total Ca^2+^ activity amplitude (Zscr.s) sites before and after exogenous application of T-peptide (100 nM) in *Cdh5-*GCaMP8 mice *in vivo*. Scale bar = 35 µm. (B) Spatiotemporal (ST) maps of all individual active Ca^2+^ sites across the entire field of view before and after T-peptide exposure in *Cdh5*-GCaMP8 mice *in vivo*. (C) T-peptide significantly decreases total Ca^2+^ activity (39 % reduction; 8423 ± 2284 vs 3257 ± 1130 ZUMS; n=5; *P<0.05; Unpaired t-test*) but (D) not the number of total events (164 ± 103 vs 104 ± 48 # events; n=5; *n.s.*; *Unpaired t-Test*). (E) Micrograph of cortical vasculature. The red box denotes where subsequent capillary flow from mice at baseline and after (F) T-peptide (100 nM) exposure was measured with a line scan. Scale bar = 20 µm. (G) Representative line scan kymographs from a capillary showing the density and velocity of RBCs (black shadows interdigitated with labeled plasma (TRITC; green) before and after pressure ejection of T-peptide (100 nm; 300 ms; 8 psi). Note the reduced flow velocity and lower RBC flux after T-peptide application (right panel). (H) Summary data showing significantly decreased RBC flux (34 ± 17 vs. 11 ± 8 RBCs/sec; n=7) after T-peptide exposure. *P<0.05;* Paired t-Test.

### T-peptide has endothelium-dependent vasoactive effects on cerebral arteries

We next tested the hypothesis that T-peptide would impact vasodilatory function in isolated pressurized pial arteries. Blood vessels were cannulated and allowed to develop myogenic tone at 80 mm Hg. Within 2 minutes of T-peptide (100 nM) application, vessels began to constriction. The endothelium-dependent vasodilator bradykinin was then used to test the effect of T-peptide on endogenous nitric oxide signaling. Bradykinin (10 µM) was administered before and 20 minutes after bath perfusion of T-peptide (100 nM). Time control experiments show that repeat application of bradykinin after 20 minutes produced a vasodilatory response that was not different from baseline (26 ± 8 vs. 21 ± 3 % change in diameter; n=3; n.s.). In contrast, after 20 minutes of T-peptide exposure, bradykinin-induced dilation was significantly attenuated (47 ± 31 vs. 12 ± 13 % dilation before and after T-peptide respectively; n=7; P<0.05) (Figure 2A-B). T-peptide did not alter the response to the endothelial Ca^2+^-activated K+ channels, SK_Ca_ and IK_Ca_, by NS309 (1 µM) (84 ± 19 vs. 67 ± 32 % dilation; n=8; n.s.) (Figure 2 C-D). These results show that T-peptide specifically attenuated endothelium-dependent signaling mechanism(s), but the capacity for hyperpolarization-dependent vasodilation through Ca^2+^-dependent activation of SK_Ca_ and IK_Ca_ channels remained intact. We further confirmed the effects of T-peptide were endothelial-specific, by testing the capacity of vascular smooth muscle to relax to the exogenous nitric oxide-donor spermine-NONOate in the presence of T-peptide. The concentration-response curve of spermine-NONOate (1-100 nM) was unaffected by T-peptide application (EC_50_ of control -7.5 µM vs. T-peptide -7.7 µM; n=3, n.s.) (Figure 2 E-F). This indicates that tau does not impair nitric oxide sensitivity in the vascular smooth muscle; instead, tau acts directly on the endothelial cells to disrupt endothelium-dependent vasodilatory function.

**Figure 2.**
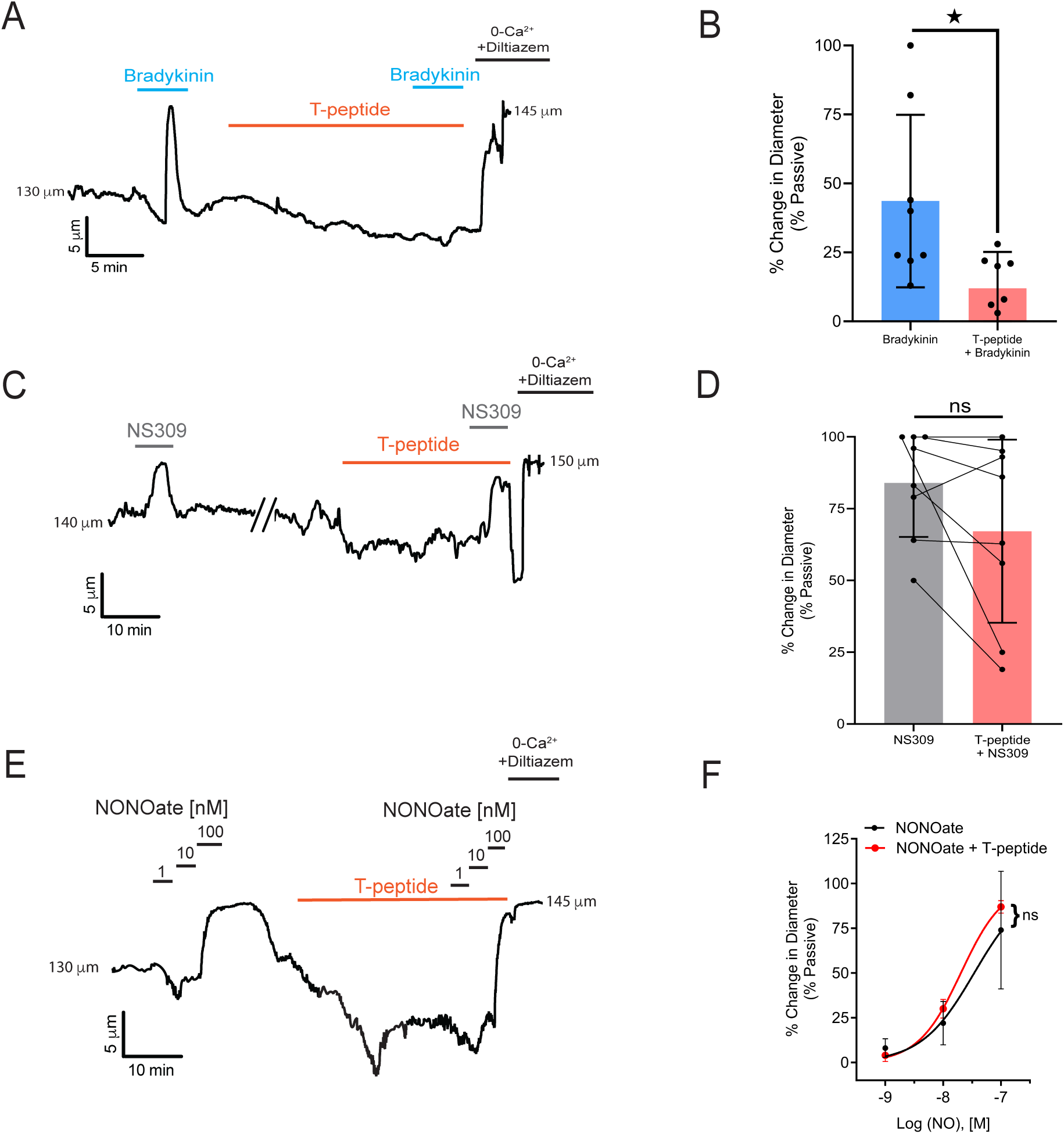
T-peptide diminishes bradykinin induced dilations but does not disrupt responsiveness to exogenous NO application. (A) Representative trace showing the response of a pressurized cerebral artery to exogenous application of bradykinin (10 µM), T-peptide (100 nM), and repeat bradykinin (10 µM) after 20 minutes of T-peptide exposure. (B) Summary data shows dilations to bradykinin (10 µM) were significantly diminished after T-peptide exposure (20 min; 100 nM) (47 ± 31 vs. 12 ± 13 % change in diameter; n=8; *P<0.05; *Unpaired t-test*). (C) Representative trace showing intact NS309 (1 µM) dilations before and after T-peptide exposure (20 min; 100 nM) in a pressurized cerebral artery. (D) Summary data shows NS309 (1 µM) dilations before and after exposure to T-peptide (100 nM) were not different (84 ± 19 vs. 67 ± 32 % change in diameter; n=8; n.s.; *Paired t-test*). (E) Representative trace showing vasodilatory responses to increasing concentrations of the nitric oxide donor spermine-NONOate (1 – 100 nM) before and after T-peptide application (20 min; 100 nM). (F) Summary data shows that T-peptide does not significantly diminish dilations to spermine-NONOate (1 – 100 nM) (LogEC_50_ of control - 7.5 ± 0.2 vs T-peptide -7.7 ± 0.04 µM; n=3; n.s; *Ordinary two-way ANOVA*).

## DISCUSSION

Tauopathy disrupts the targeted delivery of nutrients to active neurons in the brain through regional control of cerebral blood flow; this impairment in local functional hyperemia is an early manifestation of degenerative neurological diseases ^6,7,18,19^. Application of tau through a cranial window impairs local blood flow, reducing the endothelium-dependent vasodilation caused by acetylcholine ^6^. Similarly, mice with tauopathy produced through expression of aggregation-prone P301S-mutant human tau also exhibit impaired endothelium-dependent vasodilation *in vivo* ^7^. Administration of tau oligomer-specific monoclonal antibody (TOMA) rescued endothelium-dependent brain microvascular responses in this mouse model of tauopathy ^7^. Our results showing increased myogenic tone, impaired endothelium-dependent vasodilation, and reduced blood flow after t-peptide application are consistent with these prior reports. Together, these studies support a model in which tau protein accumulation in the brain leads to dysfunction of cerebral blood vessels, affecting their ability to regulate blood flow and maintain proper vascular tone. This dysfunction can impair blood flow to brain cells, limiting the delivery of oxygen and nutrients and the removal of waste products, thereby contributing to neurodegenerative processes.

Previously, the deranged cerebrovascular responses to tau were attributed to alterations in neuronal nitric oxide synthase (nNOS) and endothelial nitric oxide synthase (eNOS) function. In one study, it was shown that tau-induced dissociation of neuronal nitric oxide synthase (nNOS) from postsynaptic density 95 reduces nitric oxide production during glutamatergic synaptic activity, leading to failure of neurovascular coupling responses required for the regulation regional blood flow ^6^. In another study, it was reported that soluble pathogenic tau enters brain vascular endothelial cells and drives cellular senescence with impaired cerebrovascular function. Specifically, cultured brain microvascular endothelial cells were shown to internalize soluble tau aggregates ^7^. The misfolded forms of tau protein contained in these aggregates promoted microtubule destabilization in cultured cells, as indicated by reduced levels of acetyl-alpha-tubulin and overall microtubule density ^7^. This impaired the microtubule-dependent transport required for endothelial nitric oxide synthase (eNOS) translocation to the cell surface and subsequent nitric oxide production and induced cellular senescence. These competing models of tauopathy-induced cerebrovascular dysfunction are not mutually exclusive, however, cellular senescence would not explain the acute vasoconstriction or reduction in endothelium-dependent vasodilation we and other have observed within minutes of application of tau to brain microvessels. Instead, we propose the novel concept that tau-induced microtubule reorganization disrupts IP_3_R clustering, thereby diminishing the ability of endothelial cells to produce the Ca^2+^ signals needed for nitric oxide production and vasodilation. It has been well established that the microtubule cytoskeleton is needed for IP_3_-evoked Ca^2+^ signals in endothelial cells ^9,10^. Microtubules support the motility of IP_3_Rs, which is important because the diffusion and subsequent localization of the channel which is needed for efficient and graded recruitment of channels with increasing stimulus strength ^11^. Microtubules interact with the TxIP motif of IP_3_Rs through the microtubule end-binding protein 3 (EB3), which is critical for clustering and functional coupling between IP_3_Rs ^12^. Any effects on microtubule structure produced by soluble pathogenic tau aggregates would therefore be likely to influence IP_3_R-evoked Ca^2+^ signals. This model is supported by our results showing that when T-peptide was applied directly to the brain vasculature of mice, we observed a decrease in total cerebrovascular endothelial Ca^2+^ activity. This novel observation depended on the use of genetically encoded Ca^2+^ biosensor mice (Cdh5-GCaMP8) and also advanced spatiotemporal mapping techniques to precisely quantify the mass of individual Ca^2+^ transients in brain capillary endothelial cells. The resulting statistical z scores (zscr·μm·s) encompass the pixel intensity across the entire spatial area of an event over time. This provides reliable quantification of event amplitude for detection of even minute fluorescence changes with high reliability. ^20^ Of note, T-peptide decreased the Ca^2+^ output generated from individual active sites without reducing the number of active sites themselves, suggesting a mechanism by which T-peptide disrupts microtubule-dependent IP_3_R clustering needed to amplify local Ca^2+^ release events. This decrease in Ca^2+^ activity was associated with a significant reduction in local cerebral blood flow.

Our findings are important because in Alzheimer’s disease (AD), Chronic Traumatic Encephalopathy (CTE) and other degenerative neurological diseases, phosphorylated tau protein aggregates in cortical sulcus around small blood vessels ^1–5^. Tau is extruded to the extracellular space from damaged neurons where it oligomerizes in a process accelerated by tau phosphorylation. Pathogenic tau oligomers can transfer across synaptic clefts in brain circuits, to promote phosphorylation, misfolding and aggregation of native tau in adjacent cells through a prion-like process. This prion-like transmission through which tau proteins adopt pathogenic self-propagating conformations characteristic of prions has been well documented ^21–23^. Interestingly, it has been shown that lung microvascular endothelial cells themselves can release extracellular tau into the blood stream during bacterial pneumonia, causing cerebral tau aggregation and cognitive deficits ^24,25^. Of note, while the T-peptide portion of tau we utilized includes the C-terminal repeat region that is known to be important in tau aggregation, recent reports have demonstrated that mutations in the N-terminal region also impact the ability of tau to form larger-order complexes ^26^. These findings warrant further effort to elucidate the potential vasoactive effects of tau protein in its full-length form, and also, in the presence and absence of post-translational modifications such as hyper-phosphorylation and acetylation that play a role in neurodegeneration ^27^.

In conclusion, our results support a novel paradigm in which tau accumulation contributes to abnormal cerebrovascular function through a mechanism specifically involving endothelial cells. We demonstrated that T-peptide reduces endothelial Ca^2+^ signaling, causes vasoconstriction, impairs endothelial-dependent vasodilation, and disrupts local cerebral blood flow. These findings are significant because they support the strategy of protecting endothelial cells from tau to prevent the loss of cerebrovascular function in dementia.

## ACKNOWLEDGEMENTS

Research reported in this publication was supported by the Totman Medical Research Trust (to M.T.N.), the European Union Horizon 2020 Research and Innovation Programme (Grant Agreement 666881, SVDs@target to M.T.N.), Leducq Foundation Transatlantic Network of Excellence (Stroke-IMPaCT 19CVD01, to M.T.N.), as well as grants from the National Institute of Neurological Disorders and Stroke (NINDS) and National Institute of Aging (NIA) (K99-AG-075175 to A.M.; R01-NS-110656, RF1-NS-128963-01 to M.T.N.; and R01-NS-119971 to Nikolaus Tsoukias, subcontract to M.T.N.), the National Institute of General Medical Sciences (NIGMS) (R35-GM-144099 to K.F. and P20-GM-135007 to M.T.N.), the National Heart, Lung, and Blood Institute (NHLBI) (R35-HL-140027 to M.T.N.), and the Department of Defense/ The Henry M. Jackson Foundation for the Advancement of Military Medicine (HU001-18-2-0016 to W.L.).

## AUTHOR CONTRIBUTIONS

Conceptualization and experimental design A.M.S., K.F., A.M., M.K., G.B.; software analysis G.H., A.M.; data acquisition A.M.S., A.M., M.K., G.B., G.E.; data analysis A.M.S., A.M., G.H.; manuscript preparation and editing A.M.S., K.F., A.M., M.K., G.B., G.H., W.L., M.T.N.; project supervision and funding acquisition K.F., W.L., M.T.N.

